# The efficacy of EphA2 tyrosine phosphorylation increases with EphA2 oligomer size

**DOI:** 10.1101/2022.06.07.495003

**Authors:** Elmer Zapata-Mercado, Gabriel Biener, Daniel McKenzie, William C. Wimley, Elena B. Pasquale, Valerica Raicu, Kalina Hristova

**Affiliations:** Department of Materials Science and Engineering, Johns Hopkins University, 3400 Charles Street, Baltimore, MD 21218; Department of Physics, University of Wisconsin, Milwaukee, 3135 N. Maryland Ave, WI 53211; Department of Biological Sciences, University of Wisconsin, Milwaukee, 3209 N. Maryland Ave, WI 53211; Tulane University School of Medicine, Department of Biochemistry and Molecular Biology, New Orleans, LA; Sanford Burnham Prebys Medical Discovery Institute, 10901 North Torrey Road, La Jolla, CA 92037

## Abstract

The receptor tyrosine kinase (RTK) EphA2 is expressed in epithelial and endothelial cells and controls the assembly of cell-cell junctions. EphA2 has also been implicated in many diseases, including cancer. Unlike most RTKs, which signal predominantly as dimers, EphA2 readily forms higher order oligomers upon ligand binding. Here we investigated if a correlation exists between EphA2 signaling properties and the size of the EphA2 oligomers induced by multiple ligands, including the widely used ephrinA1-Fc ligand, the soluble monomeric m-ephrinA1, and novel engineered peptide ligands. We used Fluorescence Intensity Fluctuation (FIF) spectrometry to characterize the EphA2 oligomer populations induced by the different ligands. Interestingly, we found that different monomeric and dimeric ligands induce EphA2 oligomers with widely different size distributions. Comparison of FIF brightness distribution parameters and EphA2 signaling parameters reveals that the efficacy of EphA2 phosphorylation on tyrosine 588, which is indicative of receptor activation, correlates with EphA2 mean oligomer size. However, other characteristics, such as the efficacy of AKT inhibition and ligand bias coefficients, appear to be independent of EphA2 oligomer size. This work highlights the utility of FIF in RTK signaling research and demonstrates a quantitative correlation between the architecture of EphA2 signaling complexes and signaling features.

## Introduction

The EphA2 receptor is highly expressed in epithelial and endothelial cells, where it triggers diverse downstream signaling pathways that control the assembly of cell-cell junctions. This receptor has been implicated in many physiological and disease processes such as cancer ^1–3^, pathological angiogenesis ^4–8^, inflammation ^4, 9–11^, cataracts^12–15^, psoriasis ^16^, and parasite infections ^2, 17^. In many cases, ligand-induced EphA2 signaling has been recognized as anti-oncogenic, and thus agents that activate EphA2 could be useful as cancer therapeutics ^18^.

EphA2 belongs to the RTK family. It is a single-pass transmembrane receptor with an extracellular region that binds the activating ligands (ephrins) and an intracellular region that contains the tyrosine kinase domain. The kinase domain is activated by autophosphorylation of tyrosine residues occurring upon close contact of neighboring EphA2 molecules. Therefore, lateral interactions of EphA2 molecules are the first required step in EphA2 signal transduction in the plasma membrane.

While most of the 58 RTKs signal mainly as dimers, EphA2, in addition, can form higher order oligomers ^19–23^. Published work has suggested that the size of the oligomers may affect signaling function. For instance, ephrinA1 immobilized on artificial lipid bilayers or nanocalipers can cause different EphA2 signaling responses depending on the size of the EphA2 oligomers induced ^24–25^. However, the exact functional dependence of EphA2 signaling on the oligomerization state of the receptor is unknown. Challenges that have plagued such investigations have been (1) limited ability to control the oligomer size of EphA2 assemblies in cells and (2) limited methods to quantify heterogeneous distributions of oligomer sizes for membrane receptors. In this study, we overcome these limitations to investigate if a correlation exists between EphA2 oligomer size and signaling properties.

This work is empowered by the recent discovery of a series of small engineered peptides that bind and activate EphA2 ^22^. Given the importance of EphA2 in cell physiology and its involvement in disease, various agents have been developed to activate or inhibit/downregulate EphA2 for research and medical applications. Examples include recombinant forms of the ephrinA1 or EphA receptor extracellular regions, antibodies, peptides, small-molecule kinase inhibitors, and RNA/DNA oligonucleotides ^2, 17, 26^. Peptides hold particular promise, since they can be engineered to bind specifically to EphA2 and activate it, while the natural ephrin ligands are promiscuous and interact with multiple Eph receptors ^27–29^. The peptides used here are either monomers or constitutive dimers. They bind to the broad and shallow ephrin-binding pocket in the extracellular region, which is easily accessible on the cell surface ^28–34^.

The peptide ligands that we study here have been shown to stimulate EphA2 signaling responses with unprecedented sub-nanomolar potency and high selectivity ^22^. Interestingly, the different dimeric peptide ligands have different potencies and different efficacies, depending on their sequence and configuration ^22^. Furthermore, the peptides and the monomeric soluble form of the ephrinA1 ligand (m-ephrinA1) have been shown to induce biased signaling compared to the widely used ligand ephrinA1-Fc ^22^. In particular, these ligands can differentially modulate two EphA2 signaling responses: EphA2 autophosphorylation on tyrosine 588 (Y588, a site in the juxtamembrane segment whose phosphorylation promotes EphA2 kinase activity and activation of downstream signaling) and inhibition of AKT phosphorylation on serine 473 (S473, a site critical for AKT activation) ^35–36^.

The signaling differences may arise because the ligands stabilize different EphA2 oligomers. The dimeric peptide ligands have been engineered from monomeric precursors through N-terminal, C-terminal, or N-C terminal linkages^22^. Based on molecular modeling, we previously hypothesized that these dimeric peptides stabilize different types of EphA2 dimers, engaging different interfaces and perhaps exhibiting different signaling properties ^22^. However, we found that all the dimeric peptides induce the formation of EphA2 oligomers. Using mutagenesis of two crystallographic extracellular interfaces, the “dimerization” and the “clustering” interface ^20^, we showed that a C-terminally linked dimeric peptide induces EphA2 oligomers that utilize both interfaces ^22^. In contrast, an N-terminally linked dimeric peptide induces EphA2 oligomers that utilize the dimerization but not the clustering interface ^22^. Here, we investigate differences in the size of EphA2 oligomers that form in response to these and other peptides and to ephrin ligands.

Quantification of the oligomer size of membrane proteins has been challenging. Previously, oligomer sizes have been assessed using Förster Resonance Energy Transfer (FRET), where oligomer sizes are not measured directly but determined from model fitting to the data ^37–42^. On the other hand, fluorescence fluctuation methods offer unique opportunities to directly quantify oligomer sizes for membrane proteins^43–46^.

Recently, a method termed “fluorescence intensity fluctuations (FIF) spectrometry” was introduced, which is particularly well suited for heterogeneous populations of oligomers ^47^. FIF spectrometry calculates the molecular brightness of fluorescent protein-tagged receptors in small segments of the plasma membrane and creates a histogram of these molecular brightness values derived from thousands of such segments. The molecular brightness, defined as the ratio of the variance of the fluorescence intensity within a membrane region to the mean fluorescence intensity in this region, is known to scale with the oligomer size. Here we use FIF to characterize the oligomer size of EphA2 oligomers that form in response to ephrinA1-Fc, m-ephrinA1, three monomeric peptide ligands and three dimeric peptide ligands with different configurations ^22^.

## Results

### FIF spectrometry

We sought to assess the oligomerization state of EphA2, labeled with eYFP, using FIF spectrometry^47^. Attachment of eYFP to the C-terminus of EphA2 via a 15 amino acid (GGS)_5_ flexible linker has been previously shown to not affect EphA2 autophosphorylation^48^. Following EphA2-eYFP expression in transiently transfected HEK293T cells without ligand treatment (Figure 1A) or treated with different ligands (Figure 1B), the plasma membrane in contact with the substrate was imaged by confocal microscopy as previously described^49^. We observed that the plasma membrane exhibits homogeneous EphA2-eYFP fluorescence in the absence of ligands (Figure 1A). However, upon ligand addition heterogeneities appear within a minute or two (Figure 1B). The appearance of such “puncta” of EphA2 fluorescence in response to ligand binding has been reported in the literature^22, 50^ and used to determine whether EphA2 mutations affect receptor functionality^21^. Interestingly, the appearance of the puncta is characteristically distinct for the different ligands (Figure 1B).

**Figure 1:**
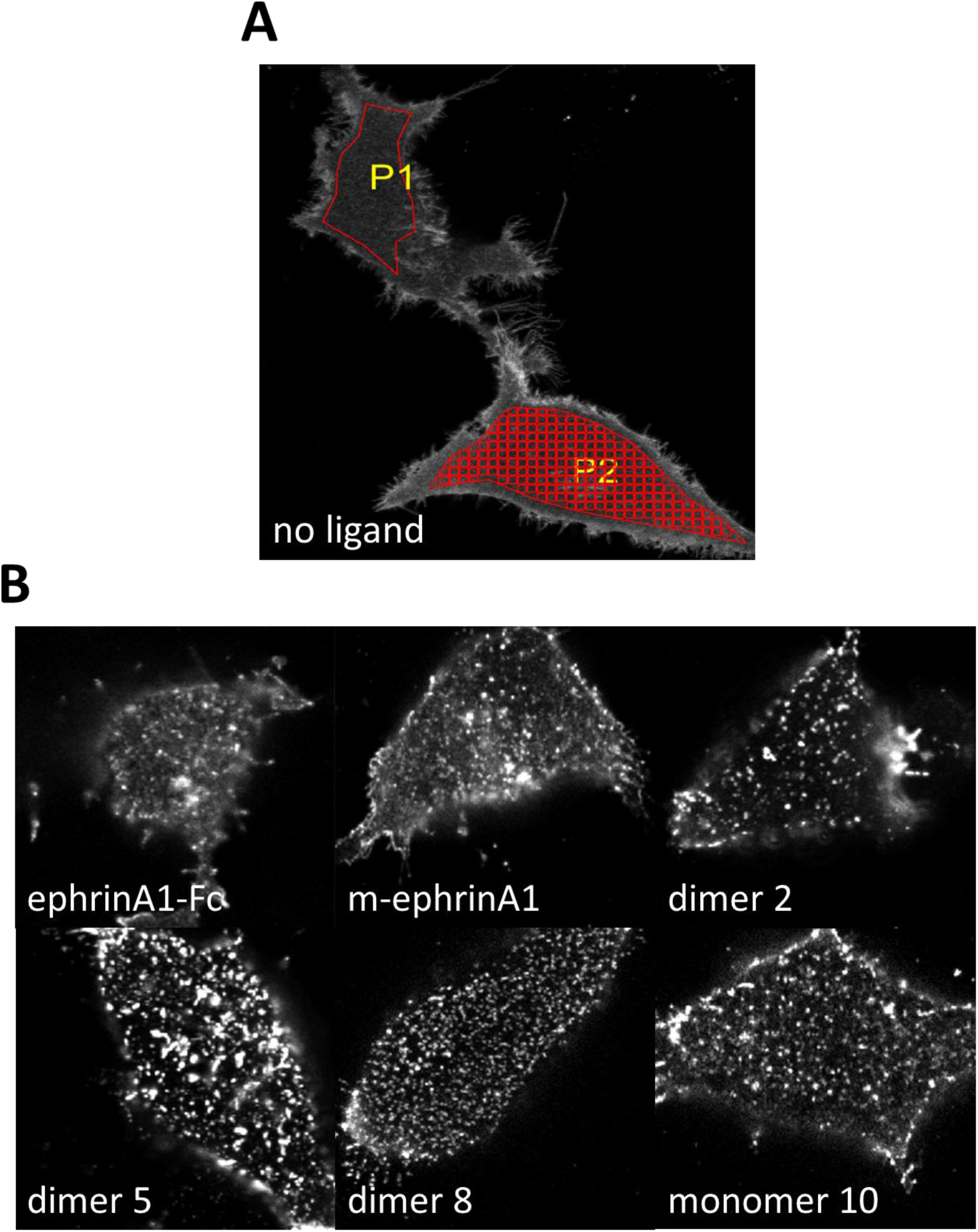
Confocal images of plasma membranes facing the solid support. HEK293T cells were transiently transfected with a plasmid encoding EphA2-eYFP. (A) No ligand stimulation. An area of the plasma membrane is selected (P1, red outline) and then segmented for FIF analysis (P2, red grid) to determine the molecular brightness in each 15 x 15 pixel segment. (B) Cells stimulated with the indicated ligands at saturating concentrations (Table 1).

**Table 1:**
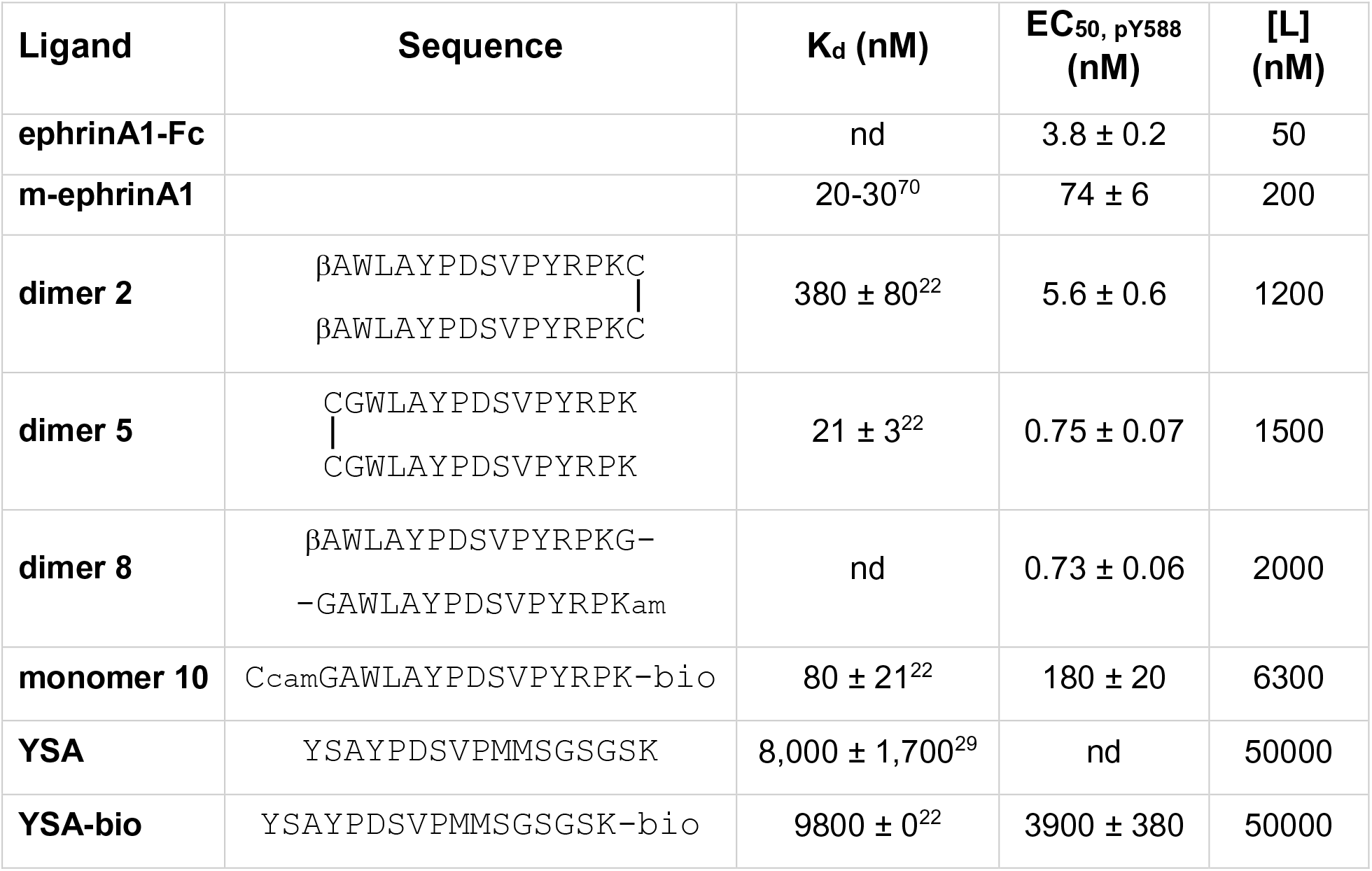
EphA2 ligands used. K_d_ is the dissociation constant as reported in ^22, 29, 69^, [L] is the ligand concentration used in the FIF experiments define EC50 and indicate the reference. nd: not determined.

Fluorescence micrographs including ∼200 to 300 cells were analyzed with the FIF spectrometry software^47^. In the first step of the analysis, a selected area of the plasma membrane (Figure 1A, P1) is divided into smaller segments with a preset size (15 x 15 pixels; Figure 1A, P2). Next, the distribution of the 225 pixel-level intensity values in each segment is fit with a Gaussian function, yielding for each segment the mean (<Ι_segment_>) and the width ((σ_segment_) of the fitted Gaussian. The variance (σ_segment_)^2^ and <Ι_segment_> are then used to calculate the molecular brightness in each segment of the plasma membrane (ε_segment_) according to equation 1 in the Materials and Methods. Finally, the brightness values from thousands of segments are histogrammed to yield molecular brightness distributions^47^.

The brightness distribution for EphA2 in the absence of ligand is shown in Figure 2A, along with the measured brightness distribution of the monomeric control LAT (Linker for Activation of T-cells)^44, 51^ and the dimeric control E-cadherin^52^. The distributions are scaled by integrating the curves and normalizing the amplitudes so that the area under the curve is the same for the three proteins. The EphA2 brightness distribution is between the brightness distributions of the monomer and dimer controls, indicating that EphA2 exists in a monomer/dimer equilibrium when a ligand is not present. This conclusion is in agreement with prior FRET studies^19, 23^.

**Figure 2:**
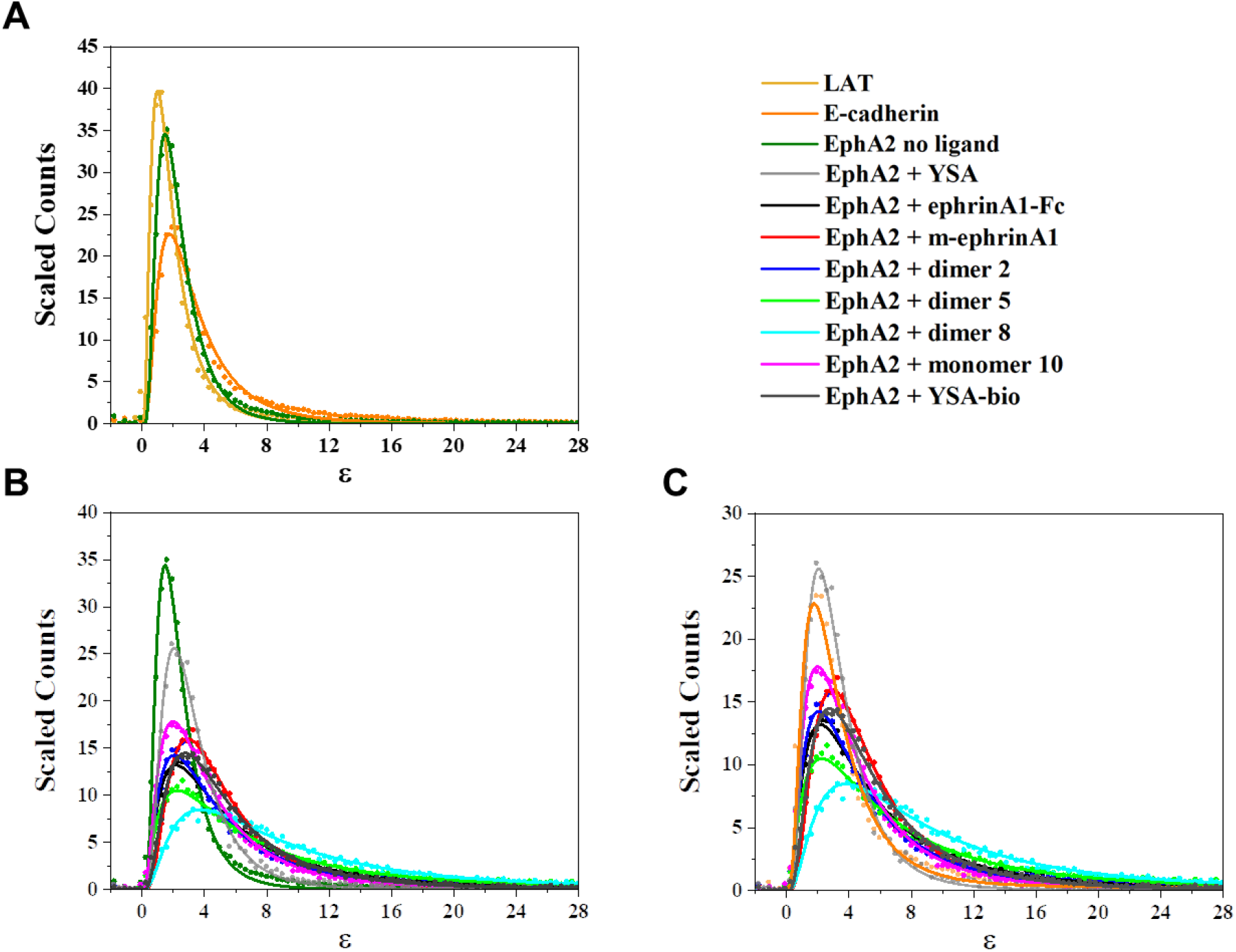
(A) Molecular brightness (ε) distributions for EphA2 in the absence of ligand, LAT (monomer control), and E-cadherin (constitutive dimer control), where all three distributions are normalized to the same area under the curve. The EphA2 molecular brightness distribution is between the monomer and dimer controls, indicating that EphA2 exists predominantly in a monomer-dimer equilibrium. (B) Molecular brightness distributions for EphA2 in the absence of ligand and in the presence of the indicated ligands, normalized to the area under the curve. The ligands shift the distributions to higher brightness. All ligands, apart from YSA, promote the formation of higher-order oligomers, while depleting the population of monomers and dimers. (C) Molecular brightness distributions for EphA2 in the absence of ligand and in the presence of the indicated ligands, compared to the brightness distribution measured for E-cadherin. The distribution in the presence of YSA is similar to the E-cadherin dimer control.

Next, we performed FIF experiments in the presence of different ligands. The ligands were used at concentrations that greatly exceed their measured dissociation constants and/or their potency (EC_50_) for EphA2 Y588 phosphorylation in cells, so that most EphA2 molecules are ligand-bound (Table 1). The brightness distributions in the presence of the ligands are compared to the brightness distribution in the absence of ligand (Figure 2B) and to the distribution of the E-cadherin dimer control^52^ (Figure 2C). All the brightness distributions were scaled so that the area under the curve is the same, allowing direct comparisons. We observed that most of the ligands shifted the distributions of brightness to higher values, indicative of the induction of higher-order oligomers. In contrast, the EphA2 brightness distribution in the presence of one of the ligands, the YSA peptide, is very similar to that of E-cadherin (Figure 2C), indicating that YSA induces the formation of EphA2 dimers rather than higher order oligomers. There are also differences among the other ligands, suggesting differences in the size of the oligomers induced. For example, the brightness distribution in response to treatment with monomer 10 is the least shifted to higher brightness values and the distribution for dimer 8 is the most shifted.

All distributions are well described by log-normal functions (see equation 2 in Materials and Methods). The two best-fit parameters of the log-normal brightness distributions, mean (µ) and standard deviation (σ), were used in equations 3 through 5 to calculate the three characteristic parameters of log-normal distributions: mean (the average brightness), median (the middle of the sorted brightness values), and mode (the position of the maximum of the distribution) (Table 2).

**Table 2:** Parameters of the molecular brightness log-normal distributions. μ and σ are the two best-fit parameters of the log-normal distributions (see equation 2). The mean, median and mode of the log-normal distributions are calculated according to equations (3), (4) and (5), respectively.

| **Whole Membrane** | **$\mu$** | **$\sigma$** | **Mean** | **Median** | **Mode** |
| --- | --- | --- | --- | --- | --- |
| **LAT** | 0.49 ± 0.02 | 0.71 ± 0.01 | 2.10 ± 0.04 | 1.63 ± 0.03 | 1.00 ± 0.03 |
| **E-cadherin** | 1.05 ± 0.02 | 0.70 ± 0.02 | 3.65 ± 0.08 | 2.85 ± 0.05 | 1.74 ± 0.05 |
| **no ligand** | 0.74 ± 0.01 | 0.60 ± 0.01 | 2.52 ± 0.02 | 2.10 ± 0.01 | 1.47 ± 0.01 |
| **ephrinA1-Fc** | 1.57 ± 0.01 | 0.89 ± 0.01 | 7.17 ± 0.09 | 4.81 ± 0.05 | 2.16 ± 0.04 |
| **m-ephrinA1** | 1.51 ± 0.01 | 0.66 ± 0.004 | 5.60 ± 0.03 | 4.51 ± 0.02 | 2.91 ± 0.02 |
| **dimer 2** | 1.45 ± 0.01 | 0.85 ± 0.01 | 6.12 ± 0.09 | 4.26 ± 0.05 | 2.06 ± 0.04 |
| **dimer 5** | 1.81 ± 0.01 | 1.00 ± 0.01 | 10.02 ± 0.16 | 6.08 ± 0.08 | 2.24 ± 0.05 |
| **dimer 8** | 2.08 ± 0.01 | 0.87 ± 0.01 | 11.66 ± 0.16 | 8.00 ± 0.09 | 3.77 ± 0.07 |
| **monomer 10** | 1.33 ± 0.01 | 0.80 ± 0.01 | 5.22 ± 0.05 | 3.78 ± 0.03 | 1.98 ± 0.02 |
| **YSA** | 1.08 ± 0.01 | 0.59 ± 0.01 | 3.52 ± 0.03 | 2.95 ± 0.02 | 2.07 ± 0.02 |
| **YSA-bio** | 1.55 ± 0.01 | 0.74 ± 0.01 | 6.20 ± 0.05 | 4.73 ± 0.03 | 2.74 ± 0.02 |

### Correlations between EphA2 signaling parameters and brightness distributions

To determine if EphA2 signaling properties correlate with the parameters of the brightness distributions, we plotted previously determined parameters that describe EphA2 signaling^22^ as a function of the mean, median, and mode of the log-normal brightness distributions (Figures 3, S1 and S2, and Table 3). The EphA2 signaling parameters we considered include (A) ligand bias coefficients (β_lig_), which describe the ability of the different ligands to inhibit AKT S473 phosphorylation as compared to increasing EphA2 Y588 phosphorylation; (B) ligand-specific efficacy of EphA2 phosphorylation on Y588 (E_top_ pY588); (C) ligand-specific efficacy of inhibition of AKT phosphorylation (E_top_ pAKT_inh_), a well-known EphA2 downstream signaling response; and (D) ligand-specific ratios of Y588 phosphorylation to AKT inhibition potencies (EC_50_ pY588/pAKT_inh_).

**Figure 3:**
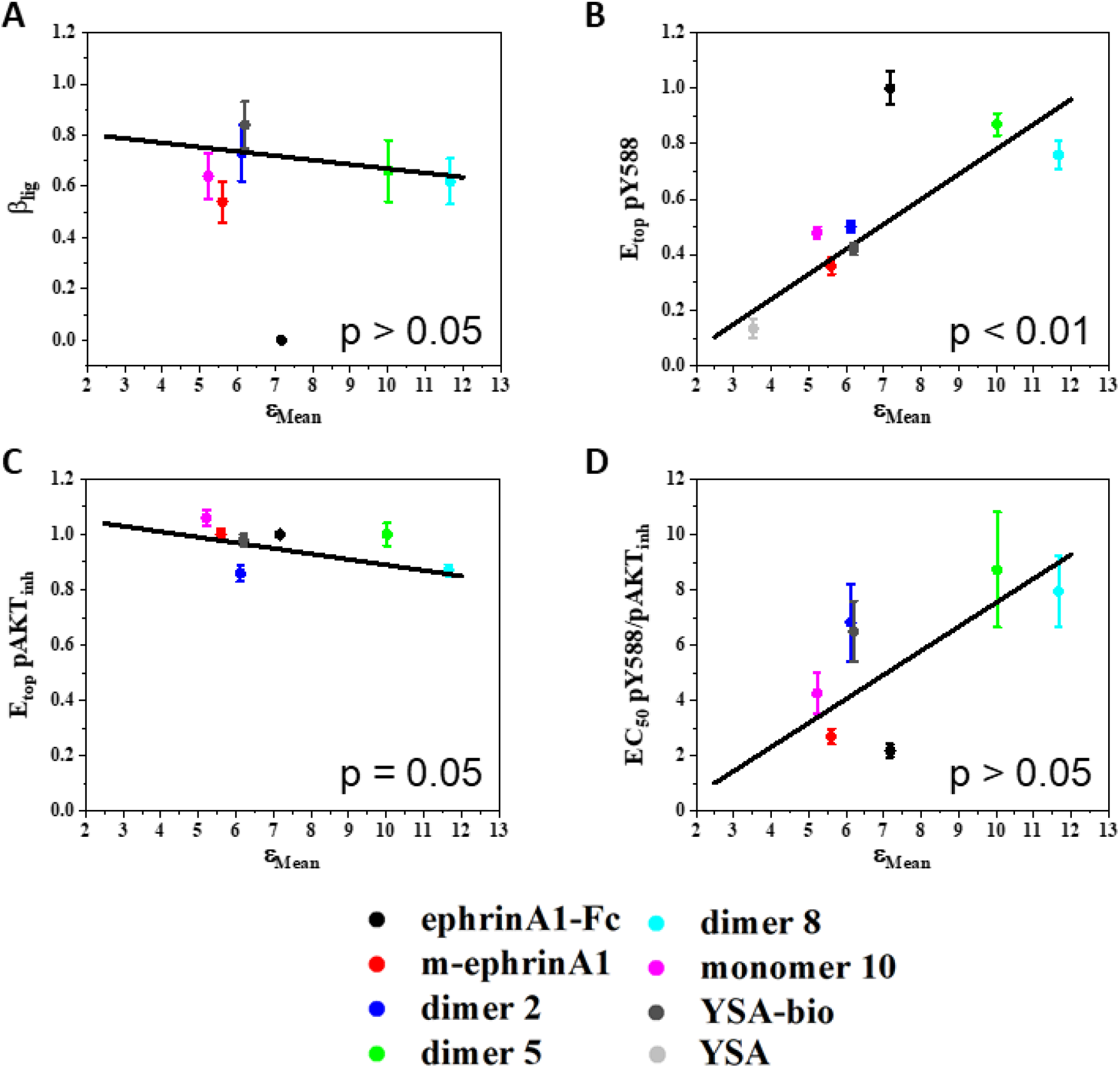
Correlation between EphA2 signaling parameters and the **mean** of the molecular brightness log-normal distributions obtained from FIF analysis of whole membranes. (A) Ligand bias coefficients versus means. (B) Ligand-specific EphA2 Y588 phosphorylation efficacies versus means. (C) Ligand-specific AKT inhibition efficacies versus means. (D) Ligand-specific ratios of Y588 phosphorylation to AKT inhibition potencies versus means. Data points: averages and standard errors from ^22^. Lines: linear fits, excluding ephrinA1-Fc. Colors are defined in Figure 2.

**Table 3:**
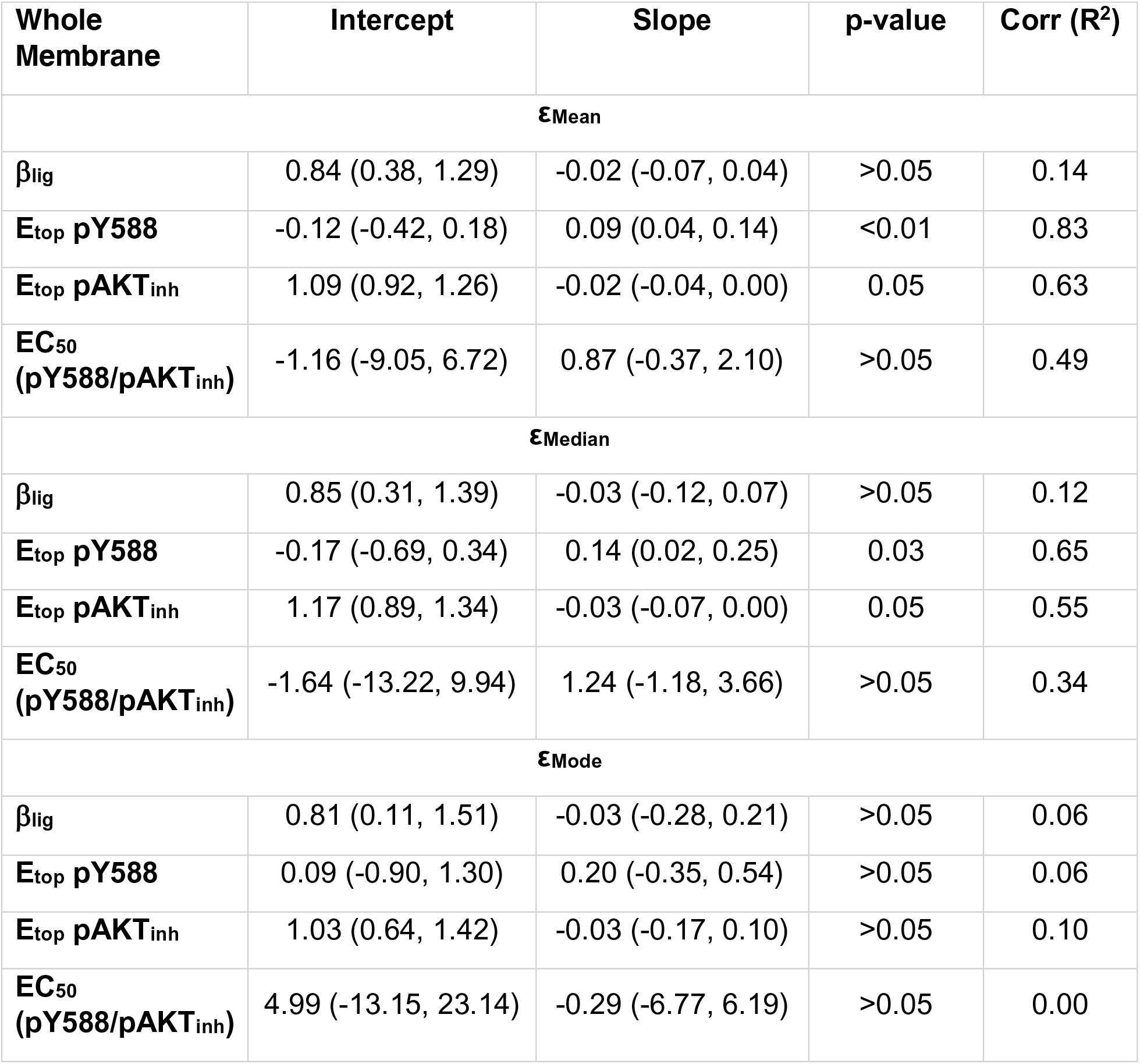
Weighted linear regression analyses to determine if there is a statistically significant correlation between the EphA2 signaling characteristics and the mean, median, and mode of the molecular brightness log-normal distributions for the whole membrane. The intercept (and 95% CI) and the slope (and 95% CI) for each linear regression are shown, as calculated from the values reported in Table 2. Correlation coefficients were determined from the fits. P-values were determined by comparing the slope to the null hypothesis of zero slope using a one sample t-test.

Calculations of bias coefficients β_lig_ for the peptide ligands and m-ephrinA1 have revealed that these ligands significantly bias signaling towards inhibiting AKT versus promoting EphA2 Y588 phosphorylation, as compared to ephrinA1-Fc ^22^. The bias coefficients for the peptide ligands and m-ephrinA1 are similar, yet the mean brightness values for these ligands are very different (Figure 3A). Furthermore, ephrinA1-Fc exhibits an intermediate brightness value. Thus, there appears to be no correlation between bias coefficients and mean brightness. This was confirmed by fitting a linear function to the peptide and m-ephrinA1 β_lig_ values (Figure 3A), and determining whether a significant correlation is present by comparing the slope to the null hypothesis of 0 slope (corresponding to no correlation) using a one sample t-test. The p-value obtained confirms that there is no correlation (Table 3). In contrast, EphA2 Y588 phosphorylation efficacy appears to increase as a function of mean brightness (Figure 3B). To determine if a correlation exists in this case, we fit a linear function to the data points for m-ephrinA1 and the peptide ligands, again excluding ephrinA1-Fc. Comparing the slope to the null hypothesis of 0 slope yielded a p value <0.01, which is indicative of a significant correlation (Table 3). Similar analyses for the efficacies of inhibition of AKT phosphorylation (Figure 3C) and the ratios of the potencies of Y588 phosphorylation to AKT inhibition (Figure 3D) show no correlation (Table 3).

Analyses for the median (Figure S1) and mode (Figure S2) of the molecular brightness distributions only revealed one additional significant correlation, which shows that the efficacy of EphA2 Y588 phosphorylation increases with the increase in median molecular brightness (Table 3).

### Molecular brightness in EphA2-eYFP puncta

To determine whether the ligand-induced increased brightness in the puncta (Figure 1B) might reflect the presence of larger EphA2 oligomers ^21^, we investigated EphA2 oligomer sizes in the puncta using FIF. Notably, FIF spectrometry can inherently filter out information about the brighter puncta because FIF data processing can ignore membrane inhomogeneities with anomalously high intensities within a segment and fit with Gaussian functions mainly the low-intensity portion of the intensity distributions^47^. This can reduce the contributions of the high-intensity pixels to the calculated mean and variance in each segment. Thus, the standard FIF analyses performed above may filter out some information about the puncta.

To specifically analyze the pixels in the puncta, we used a recent augmentation of the FIF method ^53^. In the first step of the augmented method, segments are subjected to a simple linear iterative clustering (SLIC) algorithm that identifies puncta and separates them from the cell membrane images for further analysis^53^ (Figure 4A shows an example of cell images after removal the pixels identified as belonging to puncta). Since the puncta are typically too small for reliable FIF analysis, in the second step of the augmented method the pixel content of at least four puncta with similar average intensities are combined into clusters, yielding a single molecular brightness value for each cluster. Brightness values derived from the clusters of puncta are then histogrammed in the third step and analyzed as described above for whole membranes. This algorithm has been previously used ^53^, and it has been argued that the inherent property of FIF to filter out extreme intensity values makes whole membrane analyses essentially equivalent to analyses of membranes from which puncta are removed by the algorithm.

**Figure 4:**
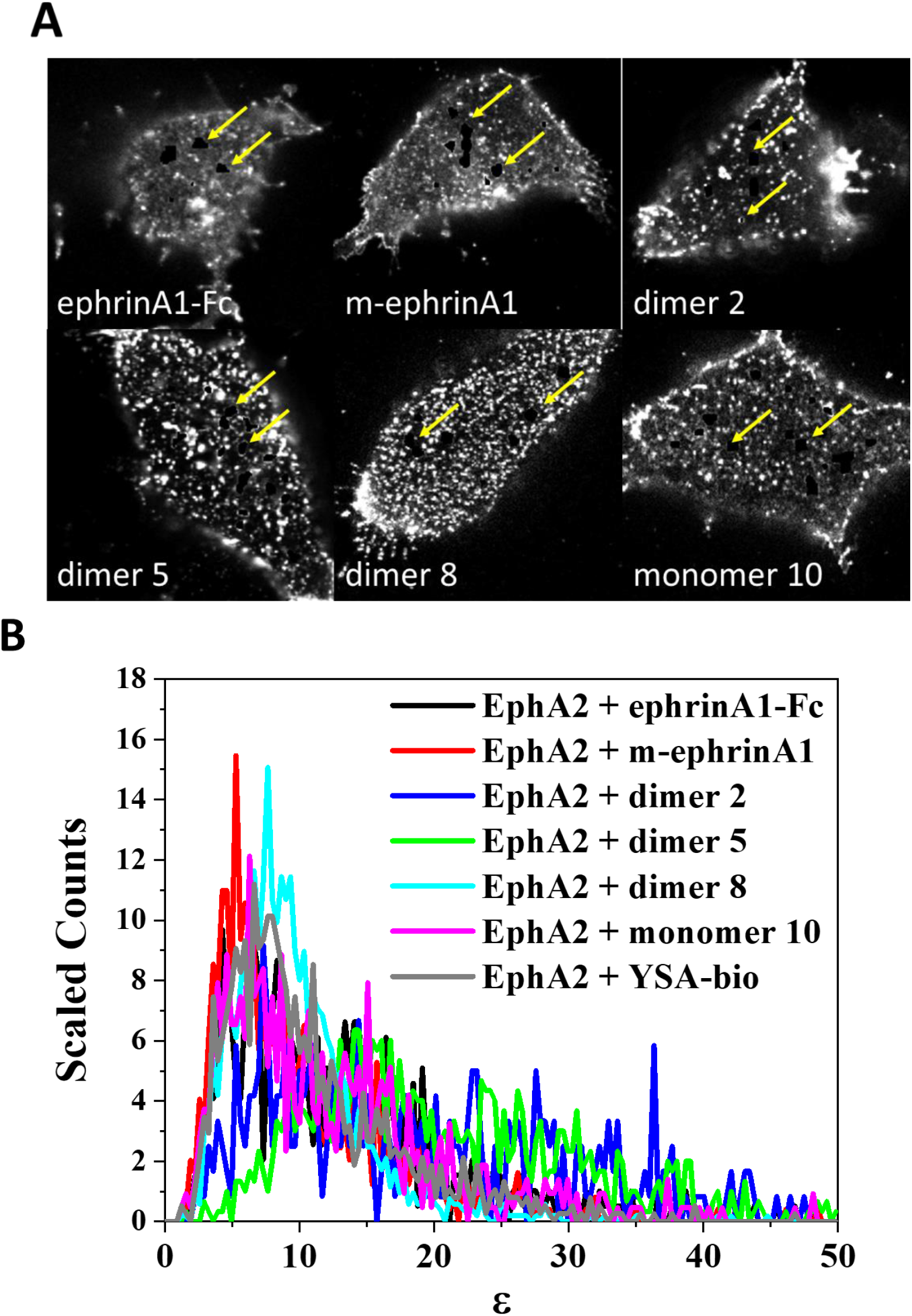
FIF analysis of EphA2-YFP puncta. (A) Cells with some puncta removed using the SLIC algorithm (pointed by yellow arrows). (B) Molecular brightness distributions for high-intensity EphA2 puncta in the presence of the indicated ligands, normalized to the same area under the curve.

Analysis of the EphA2 puncta revealed that the brightness distributions for all the ligands are shifted to higher brightness as compared to whole membranes (Figure 4B and Figure 2B). Comparison of the EphA2 concentrations obtained from the whole membrane and puncta analyses shows average concentrations ∼3 times higher in the puncta (Figure 5). However, the concentration distributions are broad, consistent with the fact that EphA2 was introduced via transient transfection.

**Figure 5:**
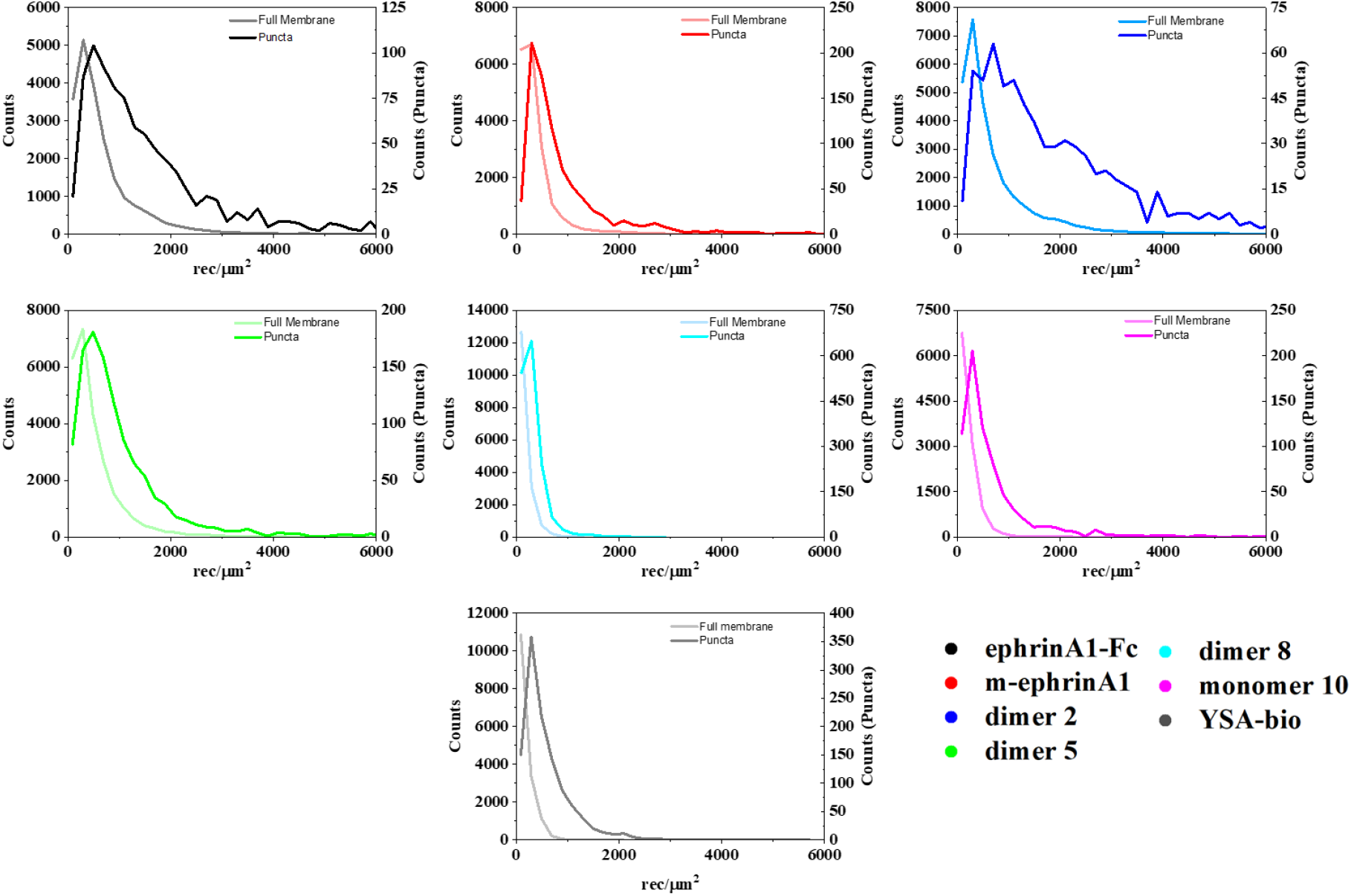
Comparison of the frequency of occurrence (counts) of EphA2-eYFP concentrations for the whole membrane (left y axis) and the high-intensity puncta (right y axis). The distributions in each panel are normalized to the same area under the curve.

Only a small fraction of the pixels were removed for puncta analysis, as shown in Figure 4A and indicated in Figure 5 by the different values of the left y axis (referring to whole membranes) and the right y axis (referring to the puncta). To directly compare the puncta brightness distributions to the whole membrane brightness distributions, we plotted them side by side while again using two different y-axis scales (Figure 6). The molecular brightness values for the puncta are shifted to the right, indicating an enrichment of higher order oligomers in the puncta. Indeed, comparison of the mean, median and mode values derived from analyses of the puncta (Table 4) with those for whole membranes (Table 2) reveals a large increase in the mode of the brightness distributions in the puncta. Curiously, the rank order of peptide mean brightness is different for puncta and whole membranes (Table S1). For example, the mean brightness ranking for dimer 8 is lower in the puncta than in the whole membrane.

**Figure 6:**
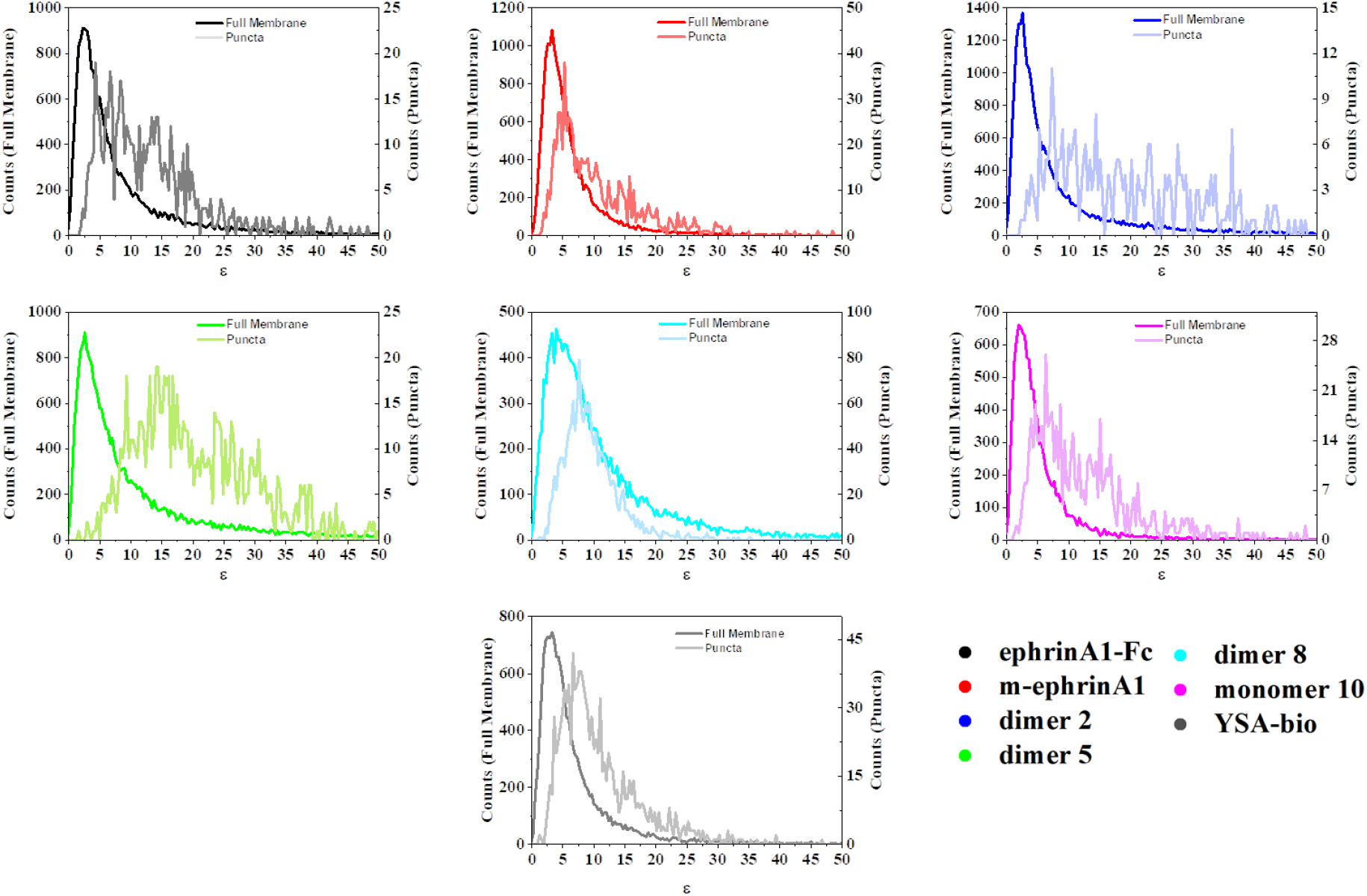
Comparison of molecular brightness distributions for the whole membrane and the puncta. The left axis refers to the brightness distributions calculated for the whole membrane. The right axis refers to the brightness distribution calculated for the puncta. The distributions derived from the puncta are shifted to higher brightness values.

**Table 4:** Mean (µ) and width (σ) of the molecular brightness log-normal distributions obtained from the high intensity puncta analysis, and the calculated mean, median, and mode for each distribution.

| **Puncta** | **$\mu$** | **$\sigma$** | **Mean** | **Median** | **Mode** |
| --- | --- | --- | --- | --- | --- |
| **ephrinA1-Fc** | $2.42 \pm 0.04$ | $0.69 \pm 0.03$ | $14.25 \pm 0.59$ | $11.21 \pm 0.40$ | $6.94 \pm 0.38$ |
| **m-ephrinA1** | $2.08 \pm 0.03$ | $0.63 \pm 0.02$ | $9.78 \pm 0.28$ | $8.03 \pm 0.20$ | $5.41 \pm 0.20$ |
| **dimer 2** | $2.91 \pm 0.08$ | $0.84 \pm 0.06$ | $26.01 \pm 2.68$ | $18.32 \pm 1.52$ | $9.09 \pm 1.22$ |
| **dimer 5** | $2.96 \pm 0.05$ | $0.54 \pm 0.02$ | $22.37 \pm 0.61$ | $19.31 \pm 0.47$ | $14.38 \pm 0.49$ |
| **dimer 8** | $2.18 \pm 0.01$ | $0.44 \pm 0.01$ | $9.76 \pm 0.10$ | $8.87 \pm 0.08$ | $7.32 \pm 0.09$ |
| **monomer 10** | $2.31 \pm 0.04$ | $0.70 \pm 0.03$ | $12.84 \pm 0.53$ | $10.07 \pm 0.35$ | $6.19 \pm 0.33$ |
| **YSA-bio** | $2.20 \pm 0.01$ | $0.56 \pm 0.01$ | $10.56 \pm 0.17$ | $9.03 \pm 0.13$ | $6.59 \pm 0.14$ |

Statistical analysis of the correlations between EphA2 signaling characteristics and mean, median, and mode of the FIF brightness distributions in the puncta did not reveal any correlations (Figures 7, Figure S3, Figure S4, and Table 5). Thus, a correlation between pY588 efficacy and mean brightness was not observed for the puncta, possibly because EphA2 signaling properties were measured using Western blotting and thus represent mean values in the whole membrane.

**Figure 7:**
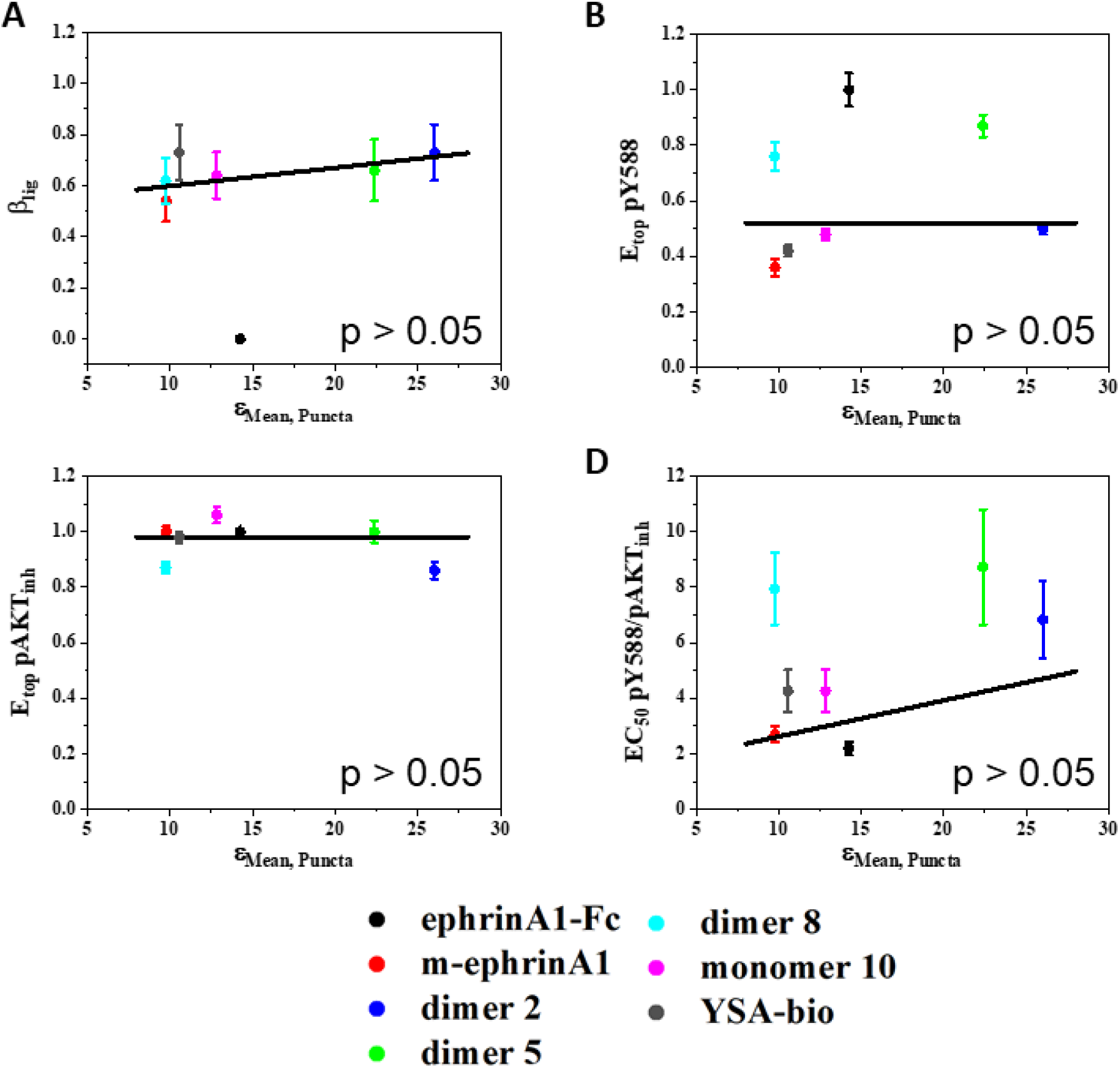
Correlation between EphA2 signaling parameters and the **mean** of the molecular brightness log-normal distributions obtained for the high-intensity puncta. (A) Ligand bias coefficients versus means. (B) Ligand-specific efficacies of EphA2 Y588 phosphorylation versus means. (C) Ligand-specific AKT inhibition efficacies versus means. (D) Ligand-specific ratios of Y588 phosphorylation to AKT inhibition potencies versus means. Lines: linear fits, excluding ephrinA1-Fc. Colors are defined in Figure 4.

**Table 5:** Weighted linear regression analyses to determine if there is a statistically significant correlation between the EphA2 signaling characteristics and the mean, median, and mode of the molecular brightness log-normal distributions for the high-intensity puncta. The intercept (and 95% CI) and the slope (and 95% CI) for each linear regression are shown for the values reported in Table 4. Correlation coefficients were determined from the fits. P-values were determined by comparing the SE_slope_ to the null hypothesis of zero slope using a one sample t-test.

| Puncta | Intercept | Slope | p-value | Corr (R<sup>2</sup>) |
| --- | --- | --- | --- | --- |
| **εMean** |  |  |  |  |
| **β<sub>lig</sub>** | 0.53 (0.38, 0.67) | 0 (0, 0.02) | >0.05 | 0.70 |
| **E<sub>top</sub> pY588** | 0.52 (-0.12, 1.16) | 0 (-0.03, 0.03) | >0.05 | 0.02 |
| **E<sub>top</sub> pAKT<sub>inh</sub>** | 0.98 (0.74, 1.21) | 0 (-0.02, 0.01) | >0.05 | 0.09 |
| **EC<sub>50</sub>**<br>**(pY588/pAKT<sub>inh</sub>)** | 1.32 (-6.78, 9.42) | 0.13 (-0.48, 0.74) | >0.05 | 0.40 |
| **εMedian** |  |  |  |  |
| **β<sub>lig</sub>** | 0.56 (0.40, 0.72) | 0 (-0.01, 0.02) | >0.05 | 0.47 |
| **E<sub>top</sub> pY588** | 0.49 (-0.15, 1.12) | 0 (-0.04, 0.05) | >0.05 | 0.04 |
| **E<sub>top</sub> pAKT<sub>inh</sub>** | 0.94 (0.74, 1.14) | 0 (-0.02, 0.01) | >0.05 | 0.01 |
| **EC<sub>50</sub>**<br>**(pY588/pAKT<sub>inh</sub>)** | 2.37 (-6.15, 10.9) | 0.06 (-0.76, 0.88) | >0.05 | 0.18 |
| **εMode** |  |  |  |  |
| **β<sub>lig</sub>** | 0.59 (0.40, 0.78) | 0.01 (-0.02, 0.03) | >0.05 | 0.24 |
| **E<sub>top</sub> pY588** | 0.19 (-0.38, 0.75) | 0.04 (-0.02, 0.10) | >0.05 | 0.60 |
| **E<sub>top</sub> pAKT<sub>inh</sub>** | 0.98 (0.67, 1.29) | 0 (-0.04, 0.02) | >0.05 | 0.13 |
| **EC<sub>50</sub>**<br>**(pY588/pAKT<sub>inh</sub>)** | -1.73 (-8.75, 5.28) | 0.89 (-0.22, 1.99) | >0.05 | 0.68 |

## Discussion

Assessment of oligomer sizes of membrane protein complexes in live cells poses unique challenges, as most methods used for soluble proteins are not applicable in the context of the native plasma membrane. Fluorescence-based methods are often the only option available to probe the oligomerization of membrane proteins suitably labeled with fluorophores. Widely used fluorescent-based techniques are FRET, fluorescence lifetime imaging (FLIM), and fluorescence fluctuation spectroscopy^51, 54–58^. Of those, the fluorescence fluctuation techniques are uniquely well suited to directly assess oligomer size, which is proportional to the molecular brightness measured^39, 47, 53^.

An example of a technique that measures fluorescence fluctuation is Number and Brightness (N&B)^43–44, 59^. N&B works by rapidly acquiring a stack of images of the same region of a cell and then computing the mean fluorescence intensity and the variance across the stack for each pixel. This yields the molecular brightness and allows for an average oligomer size to be easily calculated by normalizing the molecular brightness measured for a protein of interest to the molecular brightness of a monomer control. However, a caveat is that a large but immobile oligomer would be invisible in N&B analyses, as no fluctuations would arise over time.

While N&B monitors fluorescence fluctuations over time, other techniques such as spatial intensity distribution analysis (SPIDA) quantify fluctuations over space^60^. SPIDA works by generating histograms of pixel-integrated fluorescence intensity from a region of interest in the cell membrane to calculate an overall molecular brightness for the entire region; brightness values from several such regions are used to estimate the average size of the oligomers in the sample. In this work, we have used FIF, a space-based intensity analysis method similar to SPIDA, which yields distributions of molecular brightness derived from hundreds of small regions in hundreds of cells, as opposed to averages over statistical ensembles of cells as obtained using SPIDA ^47^. FIF spectrometry is particularly well suited for characterizing heterogeneous distributions of oligomer sizes and larger oligomers that may exhibit slow movement. It also yields concentrations of fluorophores within each image segment, which provides an additional dimension to the brightness spectrograms^47^.

Here we demonstrate that FIF spectrometry can be used to study the association of EphA2 into dimers and higher order oligomers in response to different ligands. EphA2 belongs to the RTK family, and thus its function is controlled via its oligomerization in the membrane. The formation of RTK dimers, at a minimum, is required for RTK activity, as dimerization brings two kinase molecules in close proximity so they can phosphorylate each other. Furthermore, it is known that the Eph receptors can form higher-order oligomers, similar to many other RTKs under certain conditions^61^. Although all ligands examined strongly induce EphA2 tyrosine phosphorylation and activation, with the exception of non-biotinylated YSA, surprisingly FIF experiments revealed that these ligands induce distinct brightness distributions for EphA2 in both whole membranes and in puncta.

We fond that ligands that promote different EphA2 extracellular arrangements can induce puncta with distinct appearance and different receptor oligomerization states, which might be responsible for distinct signaling properties ^22^. The YSA peptide is the only one of the ligands we examined that does not bridge two EphA2 ligand-binding domains ^22^. Our FIF experiments substantiate previous FRET experiments ^19, 62^, showing that YSA promotes the assembly of EphA2 dimers but not higher order oligomers, although the underlying mechanism remains unknown. Interestingly, YSA-induced dimerization occurs with only a small increase in receptor autophosphorylation ^22^, which conforms well with the correlation we have established between EphA2 Y588 phosphorylation and oligomer size (Figure 3B).

The fitting of the FIF distributions with log-normal functions revealed a large difference in the mode (the most frequent value) of the distributions, when comparing whole membranes to puncta (compare Tables 2 and 4). The increase in the mode ranged from 2-fold for m-ephrinA1 and monomer 10 to 6-fold for dimer 5. Thus, the most common oligomer size is higher in the puncta for all ligands. We further found that the mean/median brightness rank order is different for whole membranes and puncta, which may be due to the different appearance of the puncta induced by the various ligands, leading to different efficiencies of pixel removal for analysis. Only a small fraction of the puncta present in the membrane were identified by the SLIC protocol and used for puncta analyses, perhaps because of the modest (∼3 times) EphA2 enrichment observed in the puncta.

A RTK may be activated by different ligands, and there is great interest in developing novel biased ligands that can preferentially modulate a subset of downstream signaling responses linked to pathogenic signaling. In previous work, we analyzed dose-response curves for different EphA2 ligands to assess bias^22^. We compared two well-known EphA2 signaling responses, autophosphorylation on Y588 and downstream inhibition of AKT, in PC3 prostate cancer cells stimulated with different ligands. The bias factor, β_lig_, revealed that all the peptide ligands and m-ephrin-A1 are significantly biased towards AKT inhibition when compared to ephrin-A1 Fc. In addition, we found that the factors used to calculate β_lig_, including the efficacy and potency of the responses, differ among the ligands^22^. To determine if a correlation exists between β_lig_, efficacies or potencies and the size of EphA2 oligomers, we compared these EphA2 signaling parameters previously measured in PC3 prostate cancer cells (which have high expression of endogenous EphA2^22^) with brightness distributions measured by FIF in HEK293T cells (in which we transiently expressed EphA2 labeled with eYFP). Different responses to ligands can be acquired in different cells lines, as long as each response (in this case phosphorylation and oligomerization) is measured for all ligands under the same conditions^63–65^. This practice is common in studies of GPCRs^63^, and we use it here for EphA2 as well. Our results in Figure 3, S1 and S2 show no correlation between bias coefficients for EphA2 and the parameters of the EphA2 brightness distributions, measured by FIF.

Although bias coefficients are not different, there are quantitative differences in the features of EphA2 signaling used to calculate bias. For instance, the efficacies of the responses induced by the different ligands are significantly different from each other^22^. The efficacy is the highest possible response that can be induced by a ligand, typically at high (saturating) ligand concentrations. Here we find a significant positive correlation between the efficacy of EphA2 Y588 phosphorylation in response to m-ephrinA1 and peptide ligands, and the oligomer size. This suggests that the efficacies of EphA2 biological responses can be modulated by agents that control oligomer size. This finding sets the stage for further investigations in different cell lines to assess the general validity of our conclusions. It will be also interesting to investigate whether correlations between activity and oligomer size exist for other Eph receptors and other RTKs in general.

## Materials and Methods

### Plasmids

The EphA2-eYFP cDNA was cloned into the pcDNA3.1 (+) mammalian expression vector^19^. The eYFP fluorescent protein was attached to the C terminus of EphA2 via a flexible 15 amino acid (GGS)_5_ linker.

### Cell Culture

Human Embryonic Kidney (HEK293T) cells were cultured in Dulbecco’s modified Eagle medium (DMEM), supplemented with 10% fetal bovine serum (ThermoFisher), 1.8 g/L d-glucose, and 1.5g/L sodium bicarbonate. Cells were seeded in 35-mm glass-bottom collagen-coated dishes (MatTek’s Corporation) at a density of 2.0×10^4^ and kept in an incubator at 37°C with 5% carbon dioxide.

### Transfection

Cells were transfected with varying amounts of DNA using Lipofectamine 3000 (Invitrogen) according to the manufacturer’s recommended protocol. Twelve hours after transfection, the cells were rinsed and starved for 12 hours in phenol red- and serum-free medium containing 0.1% w/v bovine serum albumin (BSA).

### Imaging

The membranes of cells transfected with EphA2-eYFP were imaged on a Leica SP8 confocal microscope using a photon counting detector. eYFP was excited using a 488nm diode laser at 0.1% to avoid photobleaching, at a scanning speed of 20Hz. Cells were subjected to osmotic stress with a hypoosmotic medium of 25% starvation medium and 75% water. The swelling induced by the hypoosmotic medium minimizes the effect of ruffles, folds, invaginations, and other irregularities in the plasma membrane, while also preventing EphA2 endocytosis induced by ligands^66^.

Aabout 100 to 150 images were collected for each ligand, containing a total of 200 to 300 cells. One ROI per cell was selected (Figure 1A), which was divided into segments of 15×15 (225 pixels) as described^47^, yielding a total of ∼10,000 to 20,000 segments per data set for each ligand. Histograms of pixel intensities were constructed for each segment, and fitted with a Gaussian function, yielding two parameters: <*I_segment_*>, the center of the Gaussian, and *σ_segment_*, the width of the Gaussian.

The molecular brightness of each segment *ε_segment_* was calculated from:

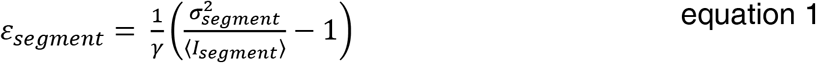

where γ is the shape factor that takes into account the beam intensity shape and the orientation of the sample relative to the beam propagation direction. Here we use a γ value of 0.5 in all cases^47^. The brightness values from thousands of segments were binned and assembled into histograms. The process of fluorescence image analysis, including ROI drawing and segmentation, concentration and brightness calculation, and further analysis was performed using a computer program described in ^67^.

The brightness distributions were fitted using OriginLab (OriginLab Corp, United States) with a log-normal function given by:

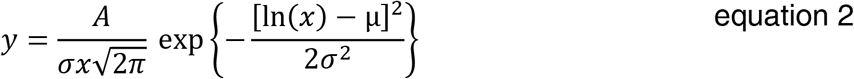

where μ is the mean of a ln(x) Gaussian distribution and σ is the width of the distribution. These two parameters were used to calculate the mean, median, and mode of the log-normal distributions according to:

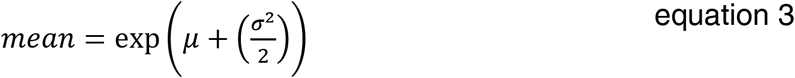

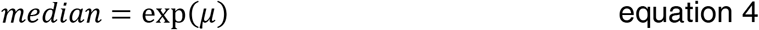

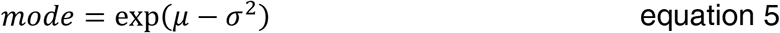

The errors of composite values were determined using propagation of errors ^68^).

To compare brightness distributions, the curves were integrated using Origin Lab, and the calculated areas were used to normalize distributions such that they have the same area.

### Correlations

GraphPad Prism 8.3 was used to assess the correlations between (i) previously reported EphA2 ligand bias coefficients, efficacies, and potency ratios^22^ and (ii) the mean, median, and mode of the molecular brightness distributions obtained from FIF. The mean, the median, or the mode were set as the independent variable (X) while the ligand bias coefficients, efficacies, or potency ratios were set as the dependent variables (Y). The data were fit to a linear function with the dependent variables weighted by 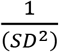, where the SD was determined from the values of SE and the total number of samples, given by the number of biological repeats N reported in ref^22^ times the different ligand concentrations used in the experiments. The slopes determined in the fits (reported in Tables 3 and 5) were compared to the null hypothesis of zero slope using a one sample t-test. P=0.05 was the cut off for the significance of the correlations.

### Puncta Identification and Analysis

Most fluorescence images of cells treated with EphA2 ligands exhibited an abundance of puncta (or “spots”), i.e., small groups of pixels with average intensities significantly higher than the surrounding membrane regions. The puncta were identified and separated for the further analysis^53^. To this end, image ROIs were subjected to a simple linear iterative clustering (SLIC) algorithm that identifies the puncta and separates them from the cell membrane images^53^. SLIC is an iterative algorithm that assigns each pixel to a certain ROI segment by calculating its “distance” to the closest segment center^69^. The distance incorporates the difference between the coordinates of the pixel and the segment center as well as the difference between the fluorescence intensities of the pixels at the two coordinates. The process is terminated when either the number of iterations reaches a chosen maximum value or the shape of the segments surrounding a punctum and the positions of the segments’ centers no longer change significantly. Full details regarding the application of SLIC to the identification of image puncta in fluorescence micrographs and subsequent analysis are provided in a recent publication ^53^. The entire protocol for puncta identification and analysis is incorporated into the program described in^67^. Practically, the process is started by segmenting an ROI using an initial segment size of 7×7, and the SLIC algorithm modifies the specific borders of the segments so that puncta of size commensurate with that of the initial segment are identified. For brightness analysis, the pixels of at least 4 puncta with similar average intensity are combined into clusters, yielding single molecular brightness values for each cluster. Brightness values derived from the clusters of puncta are then histogrammed and analyzed in a manner similar to those of whole membranes.

## Supporting information

EZM-Supplemental

