## Supplementary material for "The efficacy of EphA2 tyrosine phosphorylation increases with EphA2 oligomer size": EZM-Supplemental

| Whole Membrane | | Puncta | |
|---|---|---|---|
| | **Mean** | | **Mean** |
| **no ligand** | 2.52 ± 0.02 | **no ligand** | N/A |
| **monomer 10** | 5.22 ± 0.05 | **dimer 8** | 9.76 ± 0.10 |
| **m-ephrinA1** | 5.60 ± 0.03 | **m-ephrinA1** | 9.78 ± 0.28 |
| **dimer 2** | 6.12 ± 0.09 | **YSA-bio** | 10.56 ± 0.17 |
| **YSA-bio** | 6.20 ± 0.05 | **monomer 10** | 12.84 ± 0.53 |
| **ephrinA1-Fc** | 7.17 ± 0.09 | **ephrinA1-Fc** | 14.25 ± 0.59 |
| **dimer 5** | 10.02 ± 0.16 | **dimer 5** | 22.37 ± 0.61 |
| **dimer 8** | 11.66 ± 0.16 | **dimer 2** | 26.01 ± 2.68 |

**Table S1**: Rank order of peptide mean brightness is different for whole membranes and puncta.

### Supplemental Figures

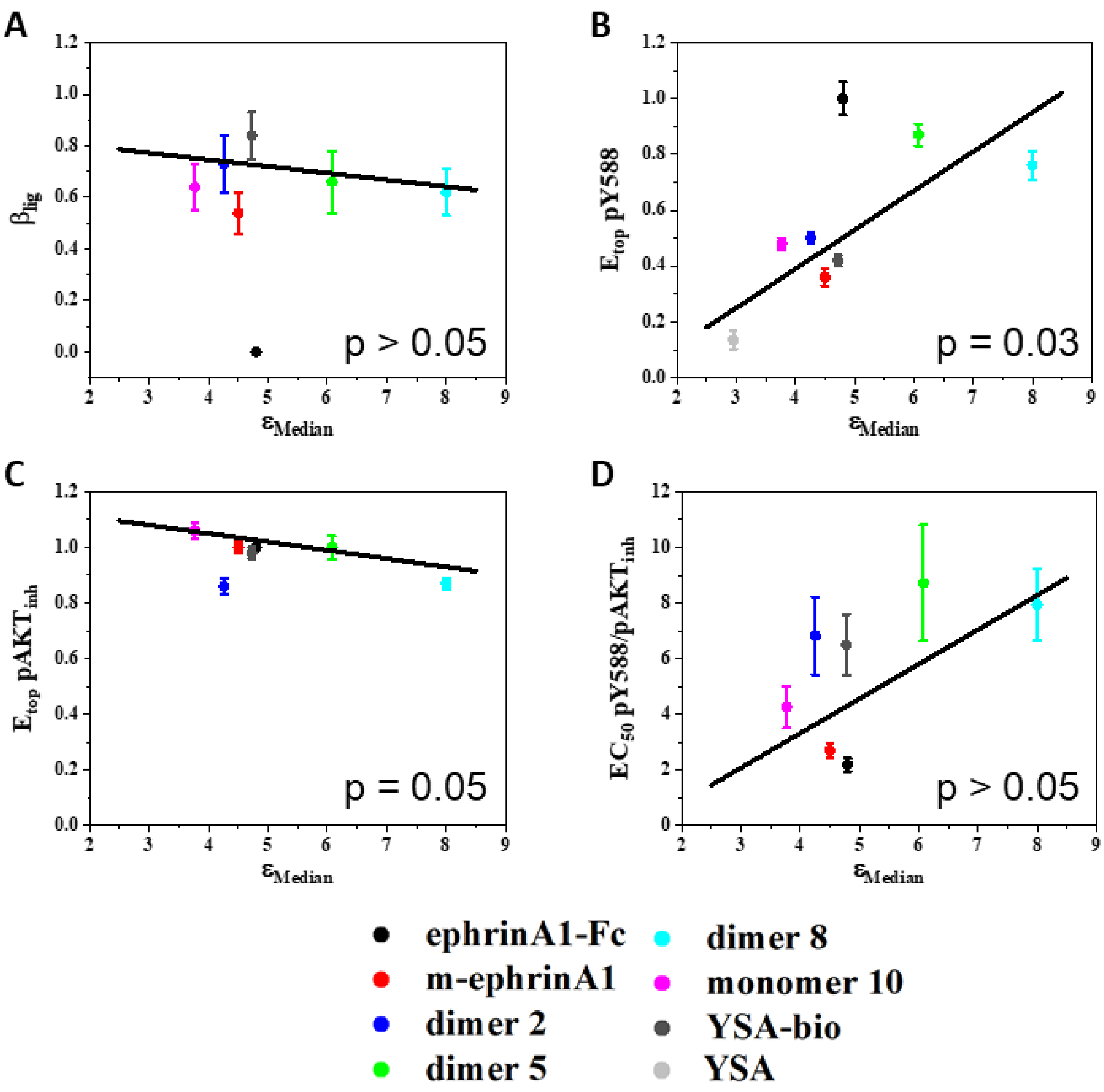


***Figure S1***: Correlation between EphA2 signaling characteristics and the **median** of the molecular brightness log-normal distributions obtained for the whole membrane analysis. (A) Ligand bias coefficients versus medians. (B) Ligand-specific efficacies of Y588 phosphorylation of EphA2 versus medians. (C) Ligand-specific efficacies of AKT inhibition versus medians. (D) Ligand-specific ratios of potencies of Y588 phosphorylation to AKT inhibition versus medians.

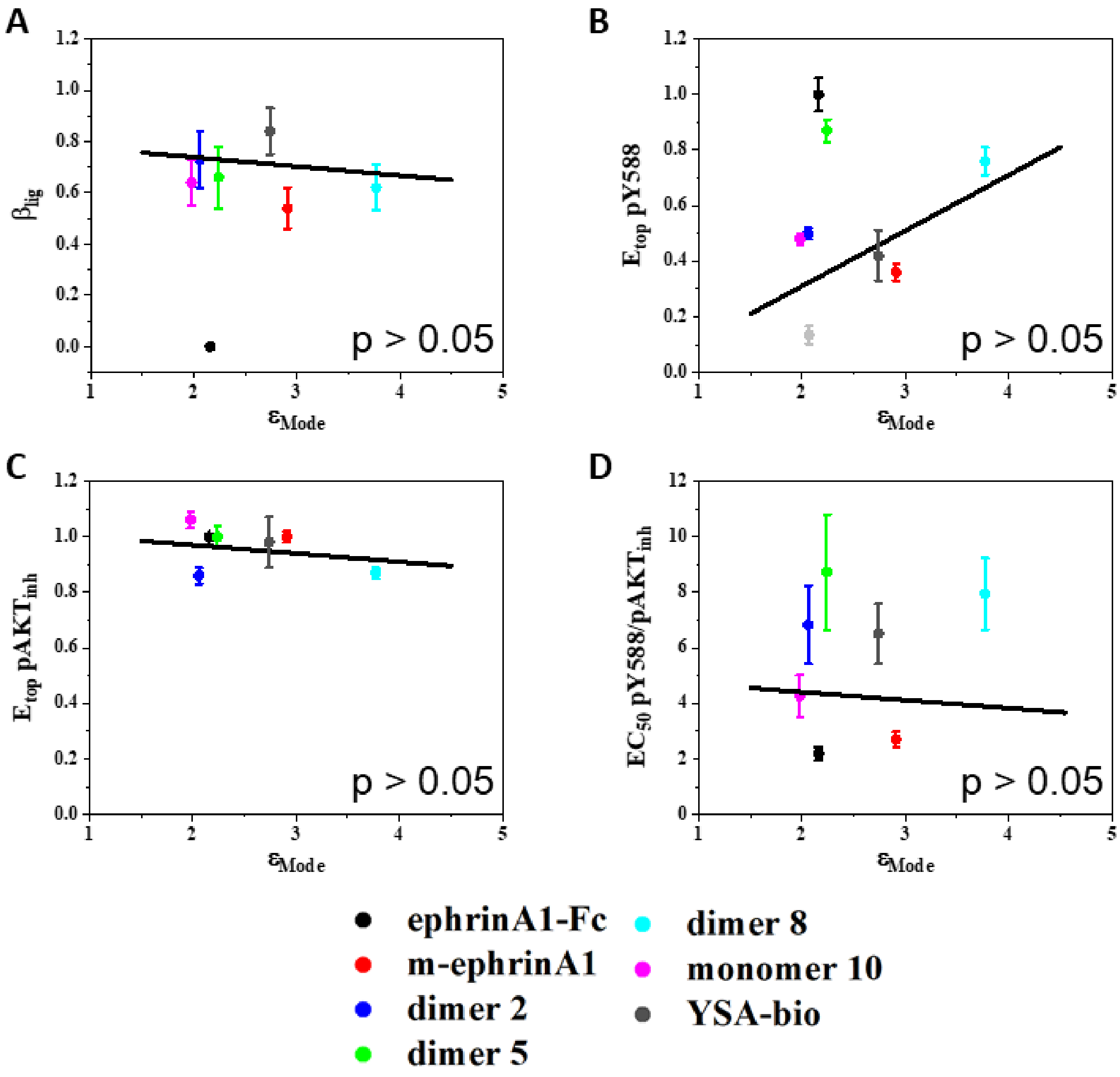


***Figure S2***: Correlation between EphA2 signaling parameters and the **mode** of the molecular brightness log-normal distributions obtained for the whole membrane analysis. (A) Ligand bias coefficients versus modes. (B) Ligand-specific efficacies of Y588 phosphorylation of EphA2 versus modes. (C) Ligand-specific efficacies of AKT inhibition versus modes. (D) Ligand-specific ratios of potencies of Y588 phosphorylation to AKT inhibition versus modes.

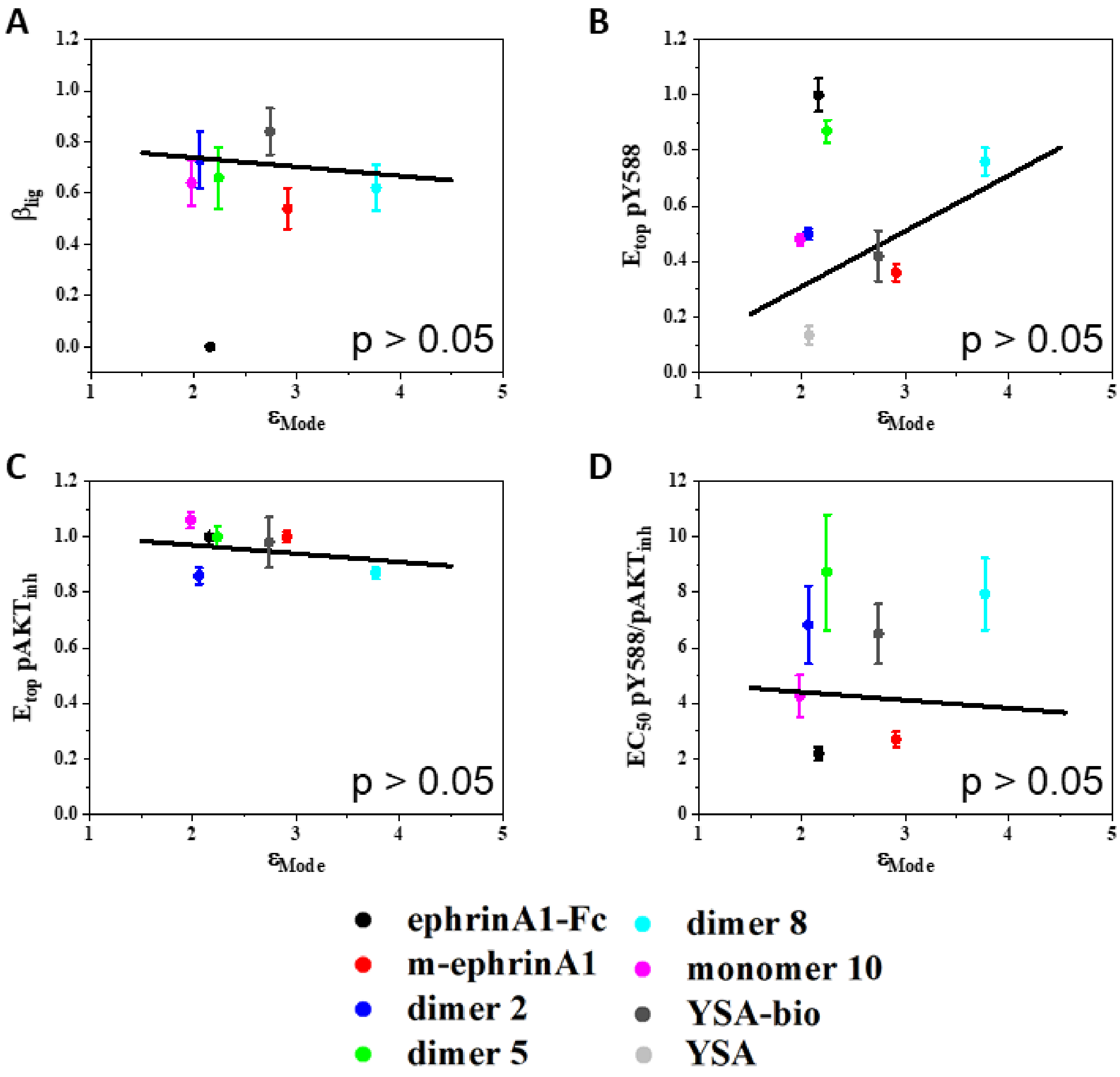


***Figure S3***: Correlation between EphA2 signaling parameters and the **median** of the molecular brightness log-normal distributions obtained for the high-intensity puncta analysis. (A) Ligand bias coefficients versus medians. (B) Ligand-specific efficacies of Y588 phosphorylation of EphA2 versus medians. (C) Ligand-specific efficacies of AKT inhibition versus medians. (D) Ligand-specific ratios of potencies of Y588 phosphorylation to AKT inhibition versus medians.

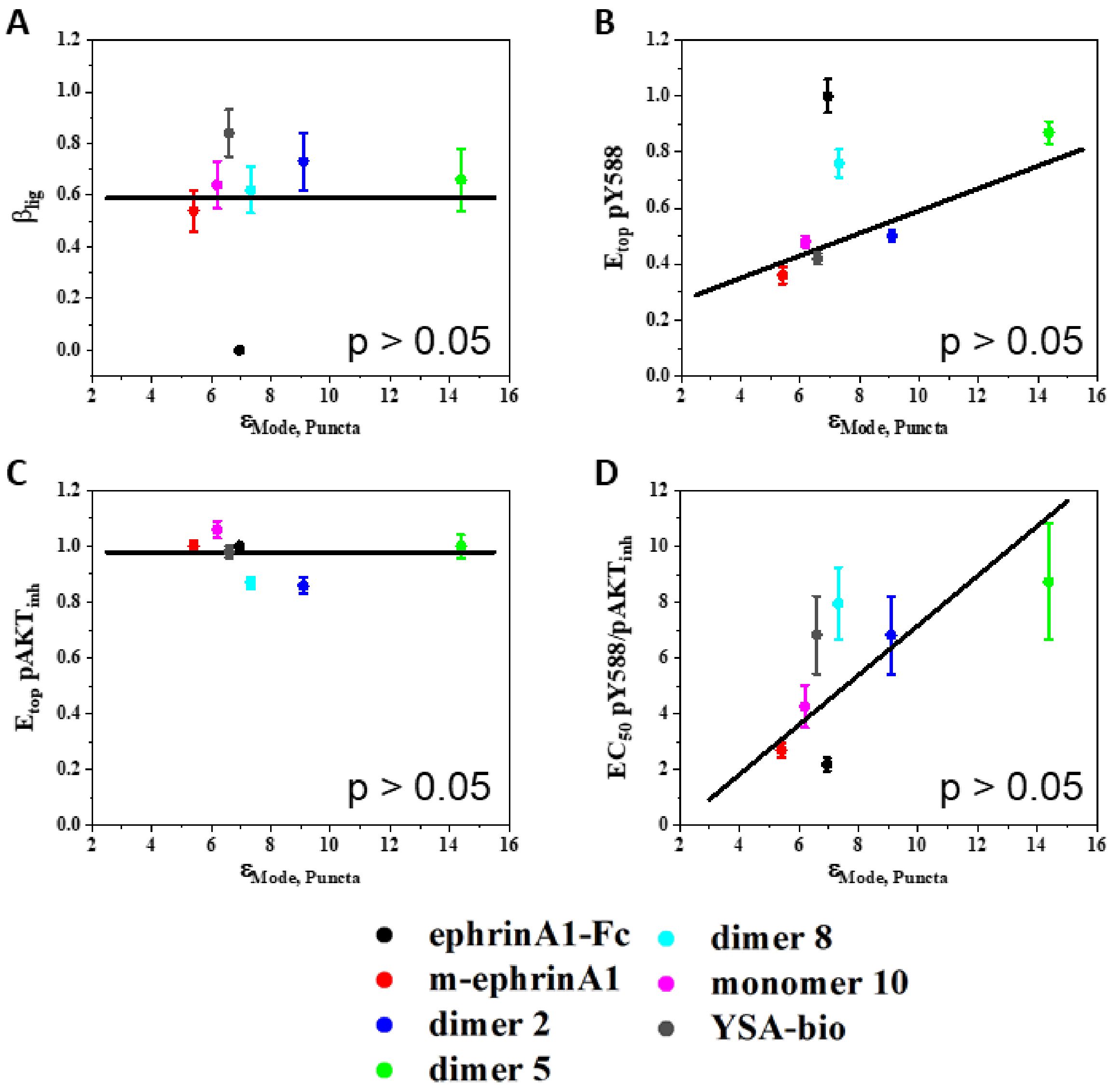


***Figure S4***: Correlation between EphA2 signaling parameters and the **mode** of the molecular brightness log-normal distributions obtained for the high-intensity puncta analysis. (A) Ligand bias coefficients versus modes. (B) Ligand-specific efficacies of Y588 phosphorylation of EphA2 versus modes. (C) Ligand-specific efficacies of AKT inhibition versus modes. (D) Ligand-specific ratios of potencies of Y588 phosphorylation to AKT inhibition versus modes.
